# “Amantadine disrupts lysosomal gene expression; potential therapy for COVID19”

**DOI:** 10.1101/2020.04.05.026187

**Authors:** Sandra P. Smieszek, Bart P Przychodzen, Mihael H Polymeropoulos

## Abstract

SARS-coronavirus 2 is the causal agent of the COVID-19 outbreak. SARS-Cov-2 entry into a cell is dependent upon binding of the viral spike (S) protein to cellular receptor and on cleavage of the spike protein by the host cell proteases such as Cathepsin L and Cathepsin B. CTSL/B are crucial elements of lysosomal pathway and both enzymes are almost exclusively located in the lysosomes.CTSL disruption offers potential for CoVID-19 therapies. The mechanisms of disruption include: decreasing expression of CTSL, direct inhibition of CTSL activity and affecting the conditions of CTSL environment (increase pH in lysosomes).

We have conducted a high throughput drug screen gene expression analysis to identify compounds that would downregulate the expression of CTSL/CTSB. One of the top significant results shown to downregulate the expression of the CTSL gene is Amantadine. Amantadine was approved by the US Food and Drug Administration in 1968 as a prophylactic agent for influenza and later for Parkinson’s disease. It is available as a generic drug..

Amantadine in addition to downregulating CTSL appears to further disrupt lysosomal pathway, hence interfering with the capacity of the virus to replicate. It acts as a lysosomotropic agent altering the CTSL functional environment. We hypothesize that Amantadine could decrease the viral load in SARS-CoV-2 positive patients and as such it may serve as a potent therapeutic decreasing the replication and infectivity of the virus likely leading to better clinical outcomes. Clinical studies will be needed to examine the therapeutic utility of amantadine in COVID-19 infection.

## Introduction

Recently a novel type of highly virulent beta-coronavirus was discovered in patients with pneumonia of unknown cause. Severe acute respiratory syndrome coronavirus (SARS-CoV-2) as detected by sequencing of the samples was found to be the cause of a severe respiratory disease in humans^1^. The outbreak of COVID-19 resulted in a global epidemic with the number of confirmed cased surpassing 722,000 in March 2020. The SARS-CoV-2 genome shares about 80% similarity with SARS-CoV and is even more similar (96%) to the bat coronavirus BatCoV RaTG13^2^. Coronaviruses are characterized by large genetic diversity and frequent recombination of their genomes, hence pose a challenge in terms of public health, currently based on 1455 viral genomes and predicted 24.5 genetic substitutions per year^3^.

Understanding the mechanism of action of the virus is a fundamental step in delineating the optimal therapeutic agents. Similar to SARS-CoV, SARS-Cov-2 enters the cell by the means of binding of cellular receptor(s) including the Angiotensin-converting enzyme 2 (ACE2) membrane bound protein^4^. ACE2 is a type I membrane protein expressed in lungs, heart, kidneys and intestine and decreased expression of ACE2 is associated with cardiovascular diseases^2,5^. The structural basis for this recognition has been recently mapped out and the cryo-EM structure of the full length viral spike protein that targets human ACE2 complex has been reported^2^. The authors show that viral S protein binds ACE2 at least 10 times more tightly, when compared to the spike protein of the previous SARS-CoV strain. The viral spike glycoprotein (S protein) mediates receptor recognition^6^. Recently the 3.5-angstrom-resolution structure of the S protein has been described^6^. The S protein is cleaved into two subunits: S1 and S2. This cleavage of S proteins by host proteases is critical for viral infection^2^ and its exist from cell via lysosomes.

Host protease dependence of SARS-CoV-2 entry is a critical step. SARS-CoV takes advantage of the endosomal cysteine proteases cathepsin B and L (CTSL and CTSB)^7,8^. Cathepsin L is a peptidase that preferentially cleaves peptide bonds with aromatic residues in P2 and hydrophobic residues in P3 position^9^. CTSL is active at pH 3-6.5, in the presence of thiol and its enzymatic stability is dependent on ionic strength^9^. Cathepsin L proteolysis was shown to be important mechanism during Ebola as well as SARS-CoV outbreak in processing viral glycoprotein before cell membrane fusion^8^. Specifically, the S protein is cleaved by host cell proteases, exposing a fusion peptide of the S2 domain. This leads to the fusion of viral and cellular membranes and the release of the viral genome into the cytoplasm of the host cell.

Cleavage of the S protein occurs between the S1 and S2 domains and subsequently within the S2 domain (S2’) proximal to the fusion peptide. Cleavage at both sites is believed to be necessary for viral entry into a host cell. The S1/S2 cleavage site of SARS-CoV-2 is between the threonine and methionine at positions 696 and 697. This S1/S2 cleavage site is identical to that of SARS-CoV which has been shown to be cleaved by cathepsin L (CatL or CTSL), a lysosomal cystein protease encoded by the CTSL1 gene. SARS-CoV-2 also has a furin-like protease cleavage site not found in SARS-CoV, between the arginine and serine at positions 685 and 686. This site may be cleaved by furin during viral egress. The S protein of SARS-CoV-2 might be also primed by TMPRSS2^4^. Inhibition of TMPRSS2 has been shown to suppress SARS-CoV infection. Furthermore (TMPRSS2), whose expression does correlate with SARS-CoV infection in the upper lobe of the lung^10^. Interfering with the spike protein processing by the host cell, whether by affecting the environment or modulating gene expression levels, hence offers a potential therapeutic strategy.

Genetic variation in CTSL gene could in theory affect the propagation capacity of the virus. Furthermore CTSL polymorphisms could affect the susceptibility to SARS-CoV-2 where for example individuals with certain genetic variant have reduced expression of Cathepsin L and in turn could be protected or have lower viral titers. Additionally, elements of hosts MHC I and CTL mediated immune responses might affect viral proliferation^11^. There are susceptibility factors spanning from ethnicity background to age related groups, to comorbid conditions^12^.

Novel therapeutics identified by high throughput screening assay shown to block the cleavage of CoV2 S (Spike) protein by Cathepsin L,Cathepsin B or TMPRSS2 (or any other protease) at predicted/selected binding sites will be a viable approach to functionally target and limit SARS-CoV-2 virus. Other therapeutic mechanisms of action could involve lowering or modulating the expression of CTSL or affecting the conditions of the CTSL lysosomal environment by modulating pH. Here we test agents that could help identify potential therapeutic agents with the capacity to decrease expression of the CTSL gene. We discuss the identification of 5 such agents and in particular Amantadine as one such compound and propose that further clinical studies should be conducted to examine whether amantadine could be useful in treating patients with COVID-19 infection.

## RESULTS

### Drug screen

To discover potential, pharmaceutical agents able to modulate molecular signatures implicated in SARS-CoV and SARS-CoV2 patho-physiology, we have screened 466 compounds belonging to 14 different therapeutic classes (Supplemental fig xx). Screening was conducted using human, retinal pigment epithelia cell line (ARPE-19) and gene expression changes were collected across 12,490 genes. The ARPE-19 cell line was initially selected as a well suited model for the study of compounds that affect neuronal type cells, in particular antipsychotics. ARPE-19 expresses a variety of well known, neuronal, cell surface receptors that include the dopamine receptor D2, the serotonin receptors 1A, 2A, and 2C, the muscarinic receptor M3, and the histamine receptor H1. Here we describe the discovery of a CTSL/B, lysosomotropic signature which might give insights into the therapeutic potential of these drugs.

We analyzed expression profile of CTSL across all 466 drugs tested. In order to find positive hits we selected only those results that caused decrease of CTSL expression by at least 33% (1.5 fold down). Top drug targets (Table1) consisted of drugs from various therapeutic areas – muscle relaxer, antihistamine, anti-epileptic, anticholirgenic and antiviral. There were no drugs that would decrease CTSL expression by more than 40%. Among the top results, Amantadine, is a known and safe antiviral agent that was previously used to treat patients with influenza A.

**Table 1.** List of top drugs affecting CTSL downregulation.

| *DrugID* | *log2(treated)*<br>*CTSL* | *log2(control)*<br>*CTSL* | *log2*<br>*difference* |
| --- | --- | --- | --- |
| *Baclofen* | 9.58 | 10.40 | -0.82 |
| *Triprolidine Hydrochloride* | 9.54 | 10.33 | -0.79 |
| *Brompheniramine Maleate* | 9.57 | 10.33 | -0.75 |
| *Amantadine Hydrochloride* | 9.62 | 10.33 | -0.70 |
| *Phenytoin* | 9.56 | 10.26 | -0.70 |
| *Atropine Sulfate* | 9.63 | 10.33 | -0.70 |

Amantadine hydrochloride (among the top drugs (within top 5 of 466) with a log2 difference is), is lysosomotropic alkalinizing agent. Physical chemical properties of amantadine leading to lysosomal accumulation^13^.. Lysosomotropic drugs affect lysosomes by pH alteration, Block of Ca^2+^ signaling, Lysosomal membrane permeabllilizations, Enzyme activity inhibition, Storage material accumulation^14^. Amantadine behaves as a lysosomotropic substance that passes easily through the lysosome membrane and accumulates in it. It could lower the pH of lysosome thus inhibit the protease activities^15^. Amantadine can also block the assembly of influenza virus during viral replication. Moreover amantadine may directly affect viral entry in direct manner down-modulating CTSL and other lysosomal pathway genes. The PK profile of the drug makes it particularly. Amantadine HCl IR is available as a 100-mg tablet (equivalent to 81 mg base amantadine) and 50 mg/5 mL syrup (equivalent to 40 mg/5 mL base amantadine), and is typically administered twice daily^16^.

Since CTSL was not the top differentially expressed transcript, we decided to extend our analysis to all the genes that were downregulated by amantadine. Among the top 500 differentially expressed probes (383 genes, all with at least 50% expression reduction) we have found 21 genes were related to lysosome (GO:005764, p=2.49×10-5) Moreover a number 1 top significant pathway by ENRICHR enrichment analysis is KEGG lysosome. Amantadine’s significant effect on lysosome pathway genes is shown on Figure 1 and Table 2.

**Table 2.**
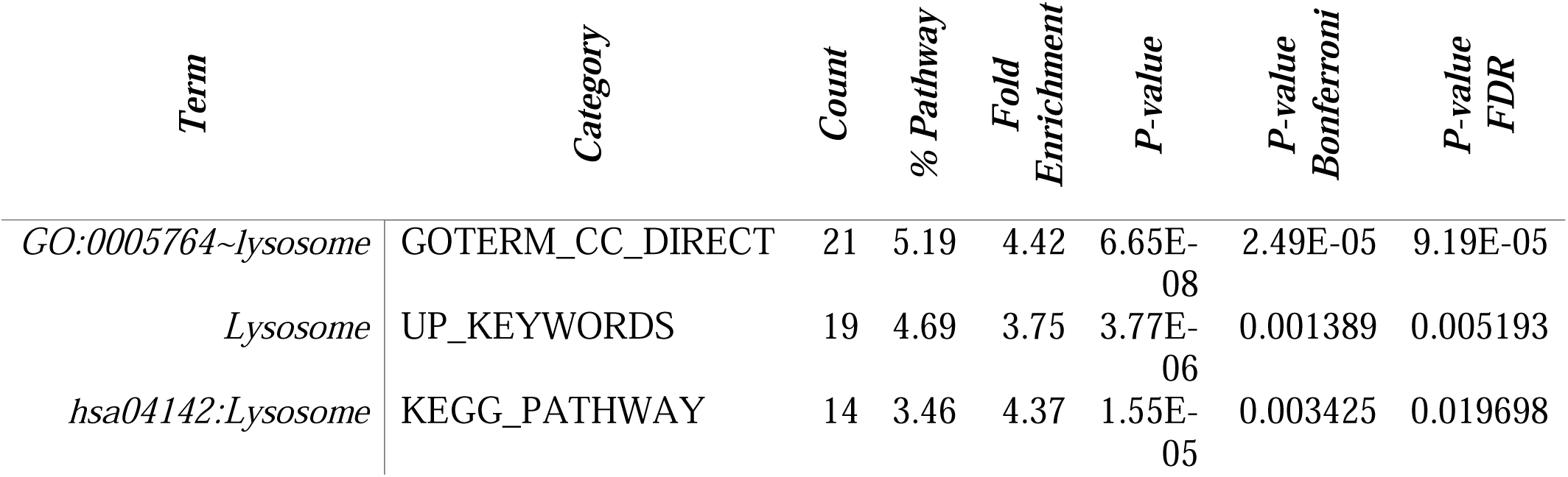

| *Term* | *Category* | *Count* | *% Pathway* | *Fold Enrichment* | *P-value* | *P-value Bonferroni* | *P-value FDR* |
| --- | --- | --- | --- | --- | --- | --- | --- |
| *GO:0005764~lysosome* | GOTERM_CC_DIRECT | 21 | 5.19 | 4.42 | 6.65E-08 | 2.49E-05 | 9.19E-05 |
| *Lysosome* | UP_KEYWORDS | 19 | 4.69 | 3.75 | 3.77E-06 | 0.001389 | 0.005193 |
| *hsa04142:Lysosome* | KEGG_PATHWAY | 14 | 3.46 | 4.37 | 1.55E-05 | 0.003425 | 0.019698 |

**Table 3.**
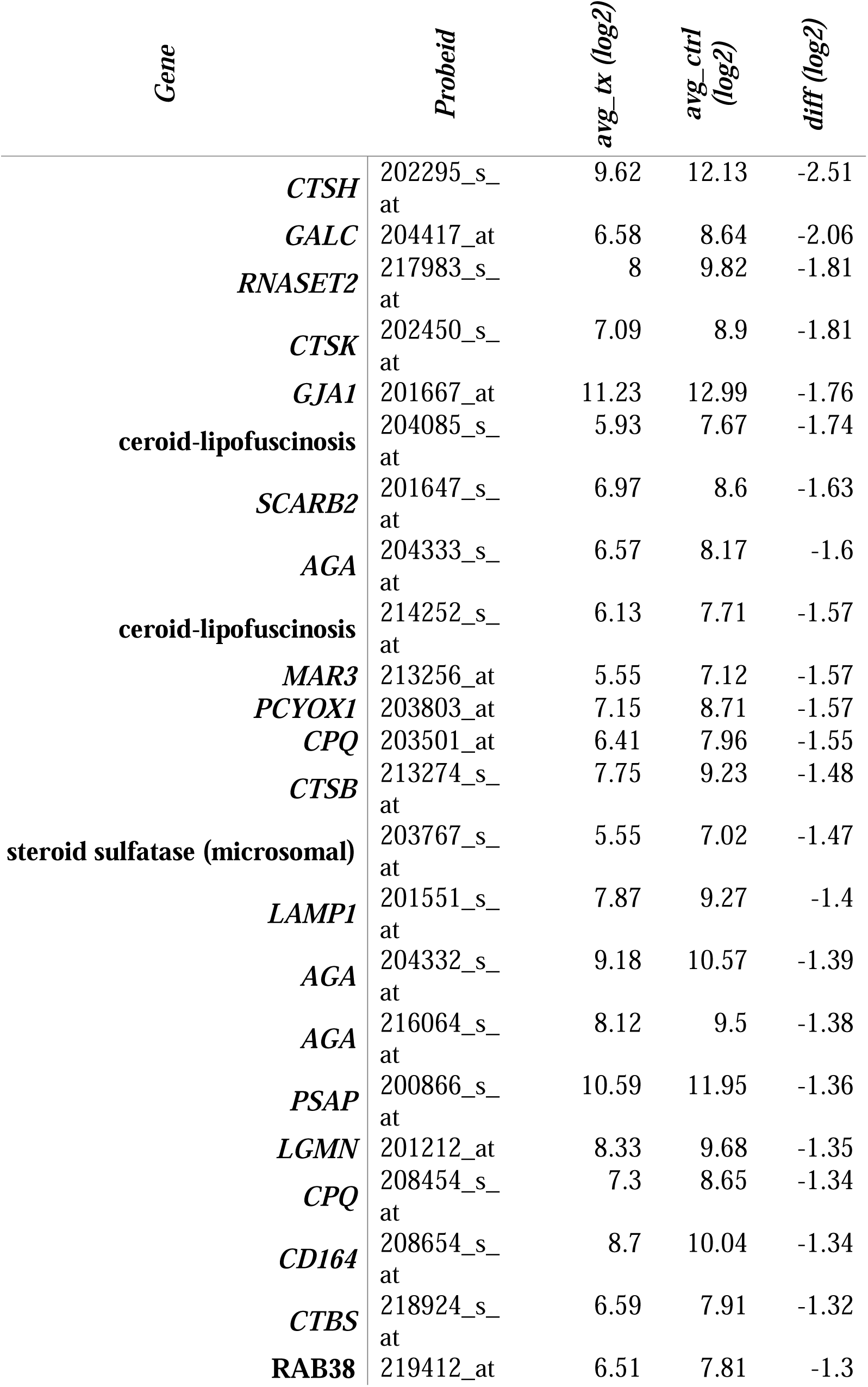
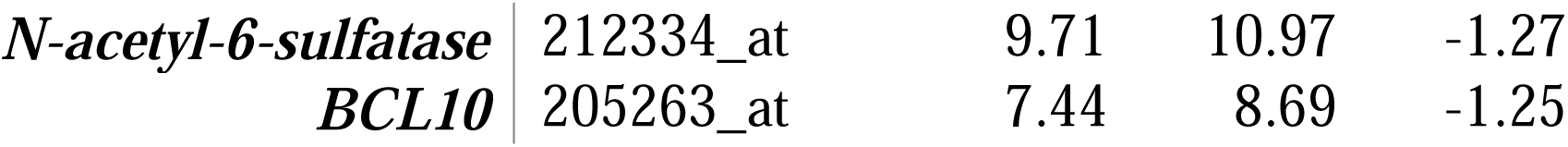
The below pathway schematic furthermore shows LAMP.

**Figure 1.**
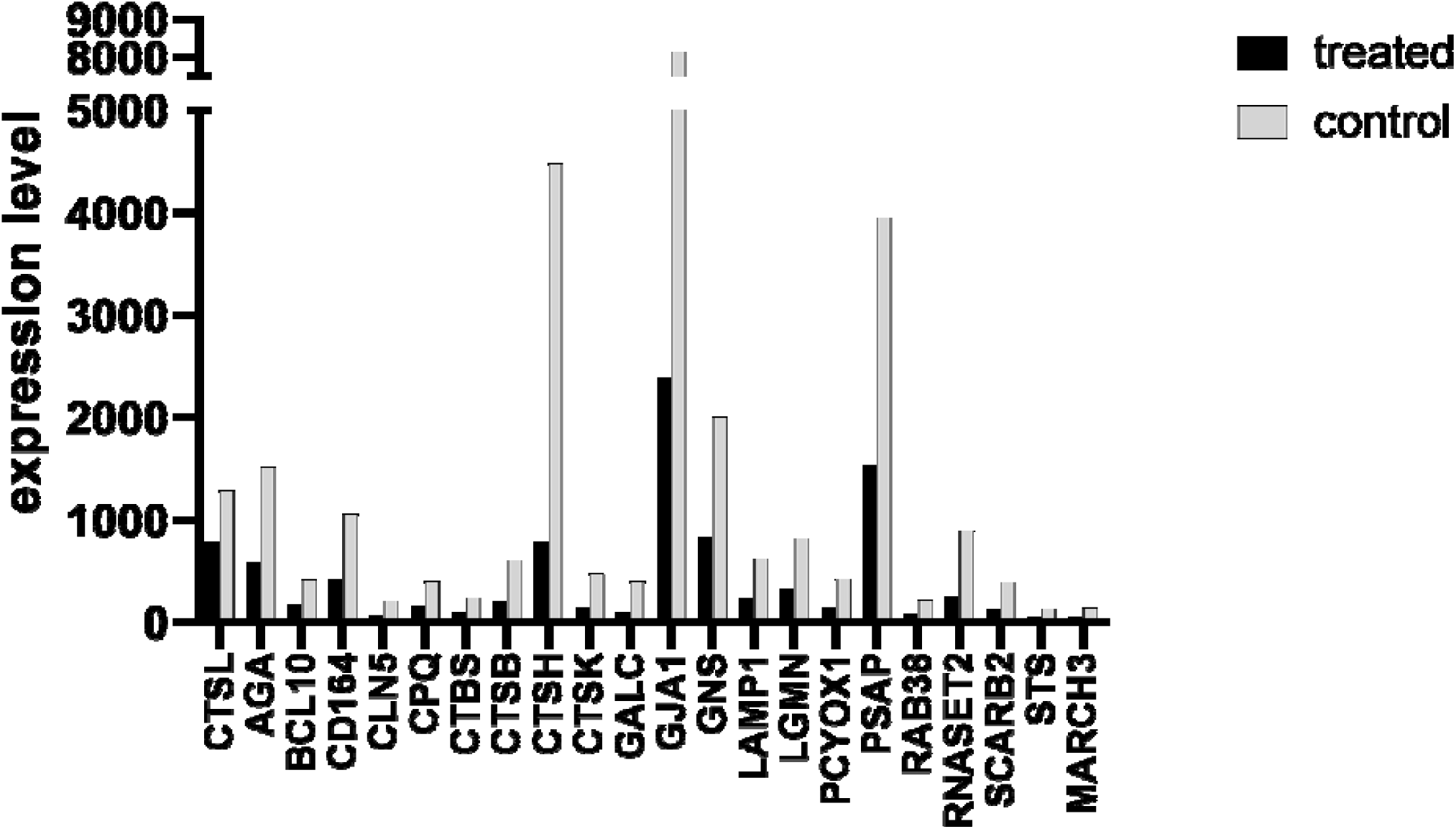
Notable difference in over-representation of genes with downregulated expression across lysosome pathway genes

We investigated natural variation of CTSL expression across ethnicities, focusing on common and rare variants. The Genotype-Tissue Expression (GTEx) project provides genotype information and gene expression levels across 49 human tissues from 838 donors, allowing us to examine the expression patterns of CTSL, both across tissues as well as across individuals. Figure 2 shows expression of CTSL across different organs and associated cell types. CTSL is widely expressed in many crucial organs (high in lungs, in nerve tibial, adipose, artery, whole blood and several others). Looking at eQTL variants in CTSL, we have found a very significant and lung specific (rs2378757) variant conferring highly variable expression. The genotype (supp fig) confers lower baseline expression and is likely associated with a better response and conversely higher expression perhaps higher viral load. We observe presence of series of splice QTLs hence variants affecting splicing ratios of transcripts such as rs114063116 significant and present in the lungs tissues. CTSL GTEx analysis points to potential protection or susceptibility of certain individuals. Interestingly the alternative splicing of the CTSL transcript in the lung further displays tissue specific regulatory programs (S.Fig1). A recent functional study points to a common variant in *CTSL* in the proximal *CTSL1* promoter, (position C-171A), confirmed to alter transcription via alteration of the xenobiotic response element^17^. This and similar other variants likely affect natural diversity in baseline expression hence viral fitness at cell entry of SARS-Cov-2. Additionally, we checked in the gnomAD^18^ database the number of rare variants and variation tolerance status of CTSL. The results show that there are on average 167 missense variants in CTSL and the gene is predicted to be loss-of-function variant tolerant with a pLI of 0.01. Together with significant eQTLs this indicates large effect of genetic variation on CTSL expression and variation thereof. TMPRSS2 is also widely expressed in multiple tissues including those in the GI system, lung, and kidney. The high expression of *CTSL* and *TRMPSS2* transcripts in a series of organs could explain the viral manifestation in these tissues such as recent studies showing viral presence in stool samples from infected individuals^19^ as well as effects of the virus seen across tissues. This is assuming that there is enough ACE2 expression on the tissue for viral S protein to bind and target the cell.

**Figure 2.**
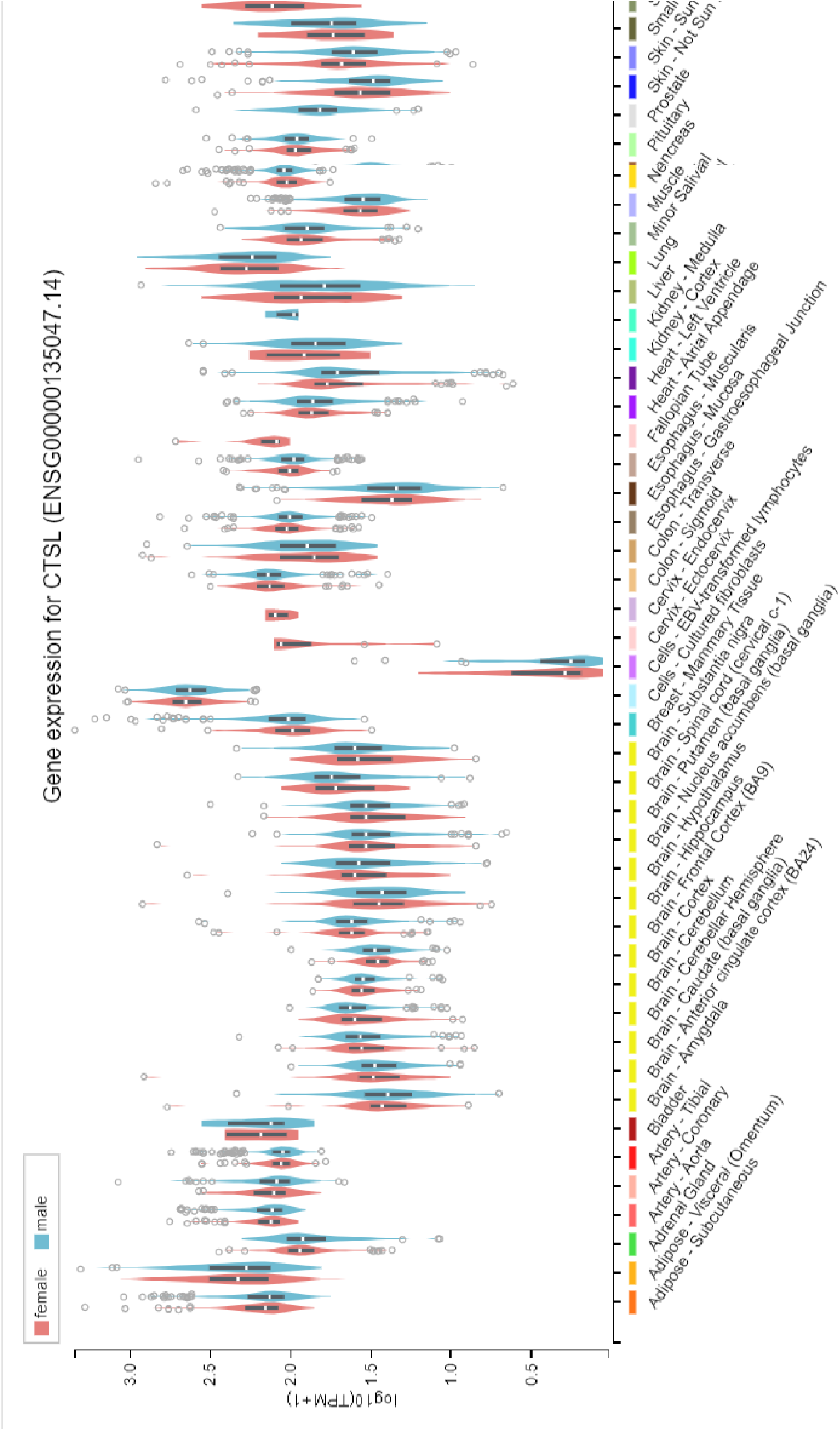
Expression of Cathepsin L (CTSL) across tissues between females (red) and males (blue)

## Discussion

Decreasing the expression of enzymes is likely a potential mechanism that would lower the capacity of the virus to exit the cell. Another viable option is lysosomal dysfunction .that can interfere in many ways with viral replication because this is where the virus is going to get assembled and get ready for subsequent replication. Given one is able to perturb the lysosomes, their function or microenvironment, you may offer protection from the virus or decrease the severity of the symptoms. Having seen that amantadine does not only down-regulate Cathepsin L, but a number of key lysosomal enzymes, we can now hypothesize that lysosomal dysfunction induced by amantadine administration can be protective from viral entry and replication. Our hypothesis that people with certain lysosomal storage diseases are resistant to one of these viruses. Along these lines there is suggestive evidence for this to be true. NPC1 (Neimman Pick type C, storage, cholesterol transport disorder) offers resistance to Ebola in patient cell lines^20, 21^. Even more interesting in the attached article, bats show selective sensitivity to Ebola versus Marburg viruses

Interfering with the lysosomal milieu can have protective effects from coronavirus which we know uses cathepsin L, a pH sensitive enzyme, to process the cleavage of the spike protein. Amantadine would be predicted by physical and chemical properties to accumulate in the lysosomes and raise a the pH, likely enough to interfere with cathepsin L function. The gene expression pattern reported in this paper suggests that a more general lysosomal program is down-regulated, likely through a common set of transcription factors. Amantadine could hence be used as a potent agent to decrease viral load if administered early enough in the course of the COVID-19 infection. Accumulation in lysosomes, If effective, could reduce viral load, may decrease intra-host organ spread and decrease associated disease severity and progression.

Further studies including clinical trials would be required in order to examine the role of amantadine administration as a treatment for COVID-19.

## Materials and Methods

### Cell culture and drug treatment

Drug screen was the same one as used in the previous study^22^. The retinal pigment epithelia cell line, ARPE-19/HPV-16, was chosen to establish a database of drug profiles because of its non-cancerous, human origin, with a normal karyotype. It can also be easily grown as monolayer in 96-well plates. Cell lines were propagated according to supplier’s specifications (ATCC Manassas, VA). Compounds were obtained from Sigma (St. Louis, MO) or Vanda Pharmaceuticals (Washington, DC). Cells were aliquoted on 96-well plates (∼2×10e5 cells/well) and incubated for 24 h prior to providing fresh media with a drug, or the drug vehicle (water, dimethyl sulfoxide, ethanol, methanol, or phosphate-buffered saline solution). Drugs were diluted 1000 fold in buffered Advanced D-MEM/F-12 culture medium (Invitrogen, Carlsbad, CA) containing nonessential amino acids and 110 mg/L sodium pyruvate. In these conditions, no significant changes of pH were expected, which was confirmed by the monitoring of the pH indicator present in the medium. A final 10 μM drug concentration was chosen because it is believed to fit in the range of physiological relevance^22^). Microscopic inspection of each well was conducted at the end of the treatment to discard any samples where cells had morphological changes consistent with apoptosis. We also verified that the drug had not precipitated in the culture medium.

### Gene expression

Cells were harvested 24 h after treatment and RNA was extracted using the RNeasy 96 protocol (Qiagen, Valencia, CA). Gene expression for 22,238 probe sets of 12,490 genes was generated with U133A2.0 microarrays following the manufacturer’s instructions (Affymetrix, Santa Clara, CA). Drugs were profiled in duplicate or triplicate, with multiple vehicle controls on each plate. A total of 708 microarrays were analyzed including 74 for the 18 antipsychotics, 499 for the other 448 compounds, and 135 for vehicle controls. The raw scan data were first converted to average difference values using MAS 5.0 (Affymetrix). The average difference values of both treatment and control data were set to a minimum threshold of 50 if below 50. For each treatment instance, all probe sets were then ranked based on their amplitude, or level of expression relative to the vehicle control (or average of controls when more than one was used). Amplitude was defined as the ratio of expression (t-v) / [(t+v) / 2] where t corresponds to treatment instance and v to vehicle instance. Each drug group profile was created using our novel Weighted Influence Model, Rank of Ranks (WIMRR) method which underscores the rank of each probe set across the entire gene expression profile rather than the specific change in expression level. WIMRR takes the average rank of each probe set across all of the members of the group and then reranks the probe sets from smallest average rank to largest average rank. A gene-set enrichment metric based on the Kolmogorov– Smirnov (KS) statistic. Specifically, for a given set of probes, the KS score gives a measure of how up (positive) or down (negative) the set of probes occurs within the profile of another treatment instance.

## Acknowledgements

We thank the investigators and patients who participated in this study.

## Supplementary Material

**S Figure 1a.**
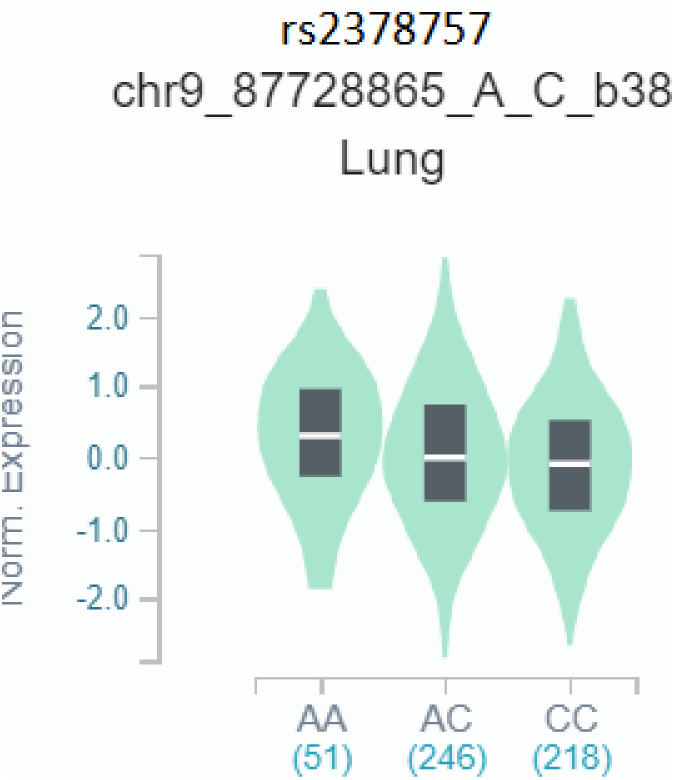

**S Figure 1b.**
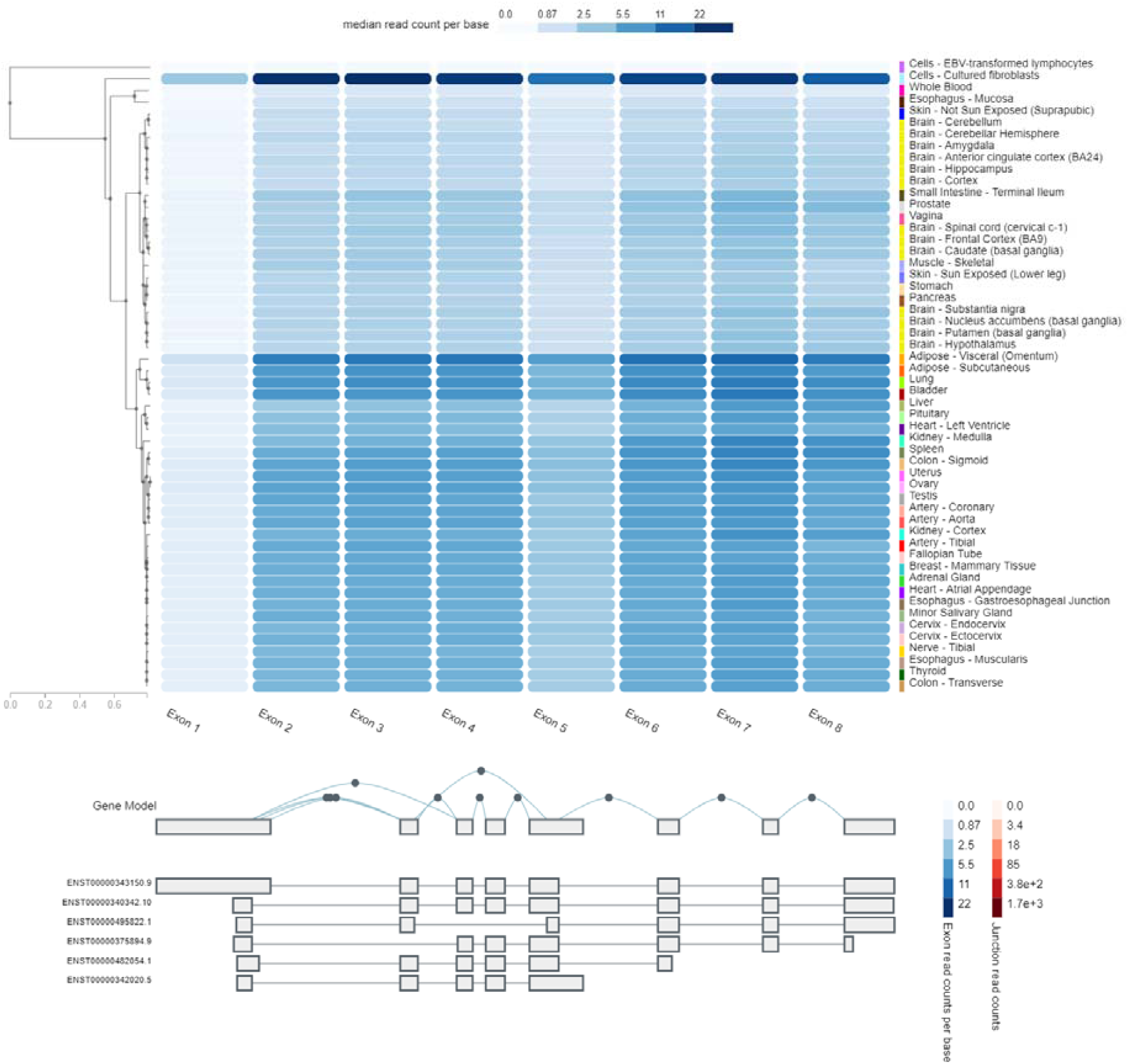
CTSL GTEX.

